# Titering of Chimeric Antigen Receptors on CAR T Cells enabled by a Microfluidic-based Dosage-Controlled Intracellular mRNA Delivery Platform

**DOI:** 10.1101/2023.03.14.532624

**Authors:** Yu-Hsi Chen, Ruoyu Jiang, Abraham P. Lee

## Abstract

Chimeric antigen receptor (CAR) T-cell therapy shows unprecedented efficacy for cancer treatment, particularly in treating patients with various blood cancers, most notably B-cell acute lymphoblastic leukemia (B-ALL). In recent years, CAR T-cell therapies are being investigated for treating other hematologic malignancies and solid tumors. Despite the remarkable success of CAR T-cell therapy, it has unexpected side effects that are potentially life threatening. Here, we demonstrate the delivery of approximately the same amount of CAR gene coding mRNA into each T cell propose an acoustic-electric microfluidic platform to manipulate cell membranes and achieve dosage control via uniform mixing, which delivers approximately the same amount of CAR genes into each T cell. We also show that CAR expression density can be titered on the surface of primary T cells under various input power conditions using the microfluidic platform.

## 1. INTRODUCTION

Chimeric antigen receptor (CAR) T-cell therapy represents a revolutionary tool for cancer immunotherapy and has significantly changed the landscape of cancer treatments. Moreover, CAR T-cell therapy has been used to effectively treat patients with relapsed and refractory B-cell acute lymphoblastic leukemia (B-ALL)^[1],[2]^ featuring two CD19-specific CAR agents (KYMRIAH and YESCARTA) approved by the Food and Drug Administration (FDA).^[3],[4]^ Despite these achievements, CAR T cells can potentially induce life-threatening toxicities, including cytokine release syndrome (CRS), neurologic toxicity, and anaphylaxis.^[5]^ CRS is one of the most common and notorious side effects of CAR T-cell therapy, followed by the infusion of CAR T cells, which triggers systemic immune activation and causes excessive inflammatory cytokine release. Recent studies have shown that the heterogeneity of CAR expression levels in cell populations determines the efficacy and safety of therapy. For instance, one study has demonstrated the influence of CAR surface density on the safety of CAR T-cell therapy; the CAR surface density can be manipulated and controlled by increasing the transduction rate with the MND promoter (myeloproliferative sarcoma virus enhancer, negative control region deleted, dl587rev primer-binding site substituted).^[6]^ Compared to the EF1α lentiviral promoter, the MND promoter has a higher packaging efficiency, but the reduced surface density of CAR molecules leads to a reduced cytokine release; therefore, it has a lower chance of triggering CRS. Moreover, another study comprehensively researched the impact of the CAR density on CAR T-cell functionality and the clinical treatment outcome.^[7]^ Phenotypic, functional, transcriptomic, and epigenomic analyses were performed to compare the high and low expressions of the CAR molecule (CAR^High^ Cell and CAR^Low^ Cell), and they concluded that CAR^High^ T cells are associated with tonic signaling and exhausted phenotypes. Hence, a precisely-controlled method that cannot only manufacture CAR T-cells efficiently but also titer the CAR expression density within an optimal range is desirable.

Currently, the most common method for manufacturing CAR T cells in a clinical setting employs viral vectors, including retroviruses and lentiviruses, owing to their high transduction efficiency.^[8]^ However, viral vectors have severe safety concerns such as insertional oncogenesis,^[9]^ transgene integration, and immunogenicity.^[10]^ Moreover, heterogeneity of CAR T-cell surface-gene expression caused by various virus copies may mitigate therapeutic efficacy or even increase the toxicity of CAR T cells.^[11]^ Bulk electroporation is another technique for delivering plasmid-based transgene systems, such as transposon/transposase systems, to introduce CAR genes into primary T cells.^[12]^ However, high voltages, ranging from tens to a few hundred volts, are required to permeabilize cell membranes and potentially cause cell mortality,^[13]^ which is a major limitation in the application of electroporation in clinical trials.^[14]^ Furthermore, several studies have shown that bulk electroporation perturbs the function of cells. DiTommaso et al. comprehensively characterized electroporation-induced disruption of gene expression, cytokine production, and the compromised in vivo biological functions.^[15]^ In addition, owing to its lack of mixing between cells and genetic materials, bulk electroporation does not offer uniform and dosage-controlled delivery across cell populations.^[16]^

To overcome the limitations of low cell viability from conventional electroporation, miniaturized microfluidic intracellular delivery systems have emerged to be a potential solution. For instance, microscale electrodes have been integrated into microfluidic platforms to achieve cell membrane permeabilization.^[17]^ The continuous-flow microfluidic electroporation platform proposed by Lissandrello et al. can achieve a high green fluorescent protein (GFP)-messenger ribonucleic acid (mRNA) transfection efficiency into primary T cells without compromising cell viability by precisely controlling the electric field exposure experienced by the cells. Another example is the utilization of a microfluidic platform to apply mechanical forces to manipulate cell membranes. Jarrel et al. demonstrated the implementation of vortex shedding in a microfluidic platform to facilitate intracellular mRNA delivery into primary human cells.^[18]^ However, these miniaturized transfection platforms cannot control the dosage of the delivered cargo or have low throughput.

In a previous study, we developed an intracellular delivery microfluidic system, the Acoustic-Electric Shear Orbiting Poration (AESOP) platform, which can disrupt the cell membrane by shearing and expanding the pores through a low-intensity electric field.^[19]^ We demonstrated high throughput (1 million cells per minute for each chip) and enabled the uniform delivery of large cargo without compromising cell viability. In this study, we explore the application of CAR T-cell manufacturing and the manipulation of CAR expression levels on primary T-cell membranes to achieve dosage control via uniform mixing and uniform shearing^[19]^. With the assistance of microstreaming vortices induced by acoustic energy, CAR mRNA is mixed uniformly with primary T cells; thus, each cell uptakes approximately the same dosage of mRNA, which leads to a similar CAR expression on the cell membrane in the cell population after mRNA translation. mRNA is utilized to transiently reprogram T cells to express CAR, which does not integrate DNA into the host genome,^[20]^ and can avoid on-target off-tumor toxicity caused by permanent CAR expression.^[21],[22]^ We show that by controlling the initial input mRNA concentration, the dosage taken by the cells and CAR expression levels can be titered. Moreover, gene expression profile and cytokine secretion function did not change after AESOP treatment. These results reveal that the delivery mechanism of AESOP can maintain the safety and function of cell therapy and improve the homogeneity of CAR T-cell products, which is a key factor in determining the outcome of a clinical treatment.

## 2. RESULTS

The schematic of isolating primary T-cells and transfecting T-cells with AESOP and electroporation, respectively, is presented in **Figure 1**. AESOP first relies on the lateral cavity acoustic transducer (LCAT) technology to trap cells inside acoustic microstreaming vortices and expose them to uniform mechanical shear. The oscillation of trapped microbubbles in lateral slanted dead-end side channels generates a first-order oscillatory flow at the air-liquid interface. The first-order oscillatory flow induces a second-order streaming flow consisting of an open microstreaming flow and a closed-loop microstreaming vortex. Open microstreaming generates bulk flow that pumps through the main channel. The LCAT is powered by a piezoelectric transducer (PZT) attached to the bottom of the device, and the acoustic energy from the PZT is transmitted to the air-liquid interfaces, causing them to oscillate and generate microstreaming vortices in the microfluidic channel. The orientation and positioning of the air-liquid cavities resulted in both bulk-flow liquid pumping and size-dependent trapping of cells. The trapped cells orbiting in these microvortices were subjected to oscillatory mechanical shear stress, with the maximum shear adjacent to the oscillating air-liquid interfaces.

**Figure 1.**
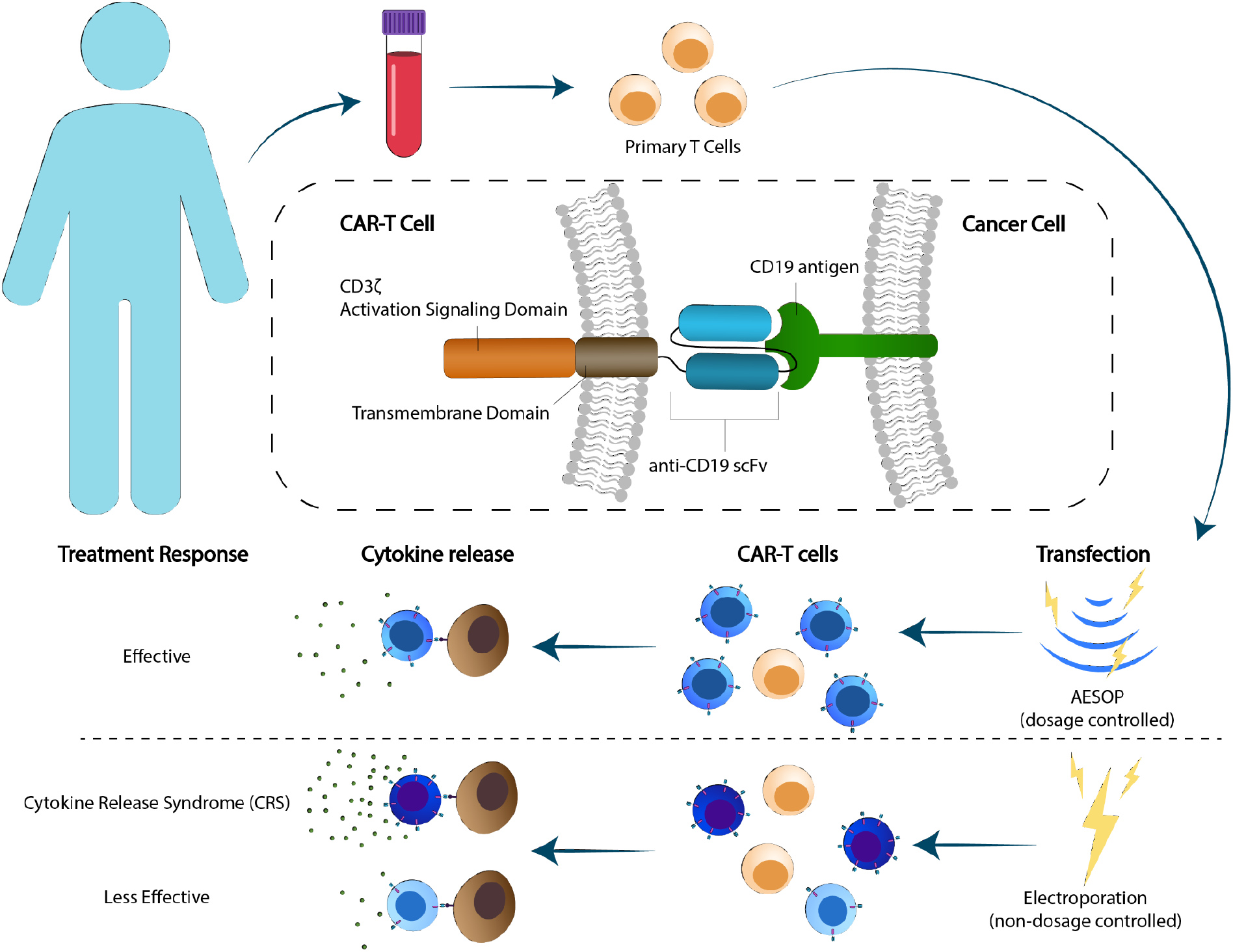
Schematic of CAR T cell generation from the extract isolation of primary T cells from healthy donors, T cell activation, and transfection with AESOP and electroporation. High, medium, and low CAR-expression levels on the T-cell membrane cause CRS as well as effective and less effective treatment responses, respectively.

### 2.1. Characterization of the shear force within acoustic microstreaming vortices

We previously used LCATs with 112 pairs of dead-end side channels combined with a single main channel to perform self-pumping, cell-cargo mixing, and cell-membrane permeabilization. We demonstrated throughput of 1 million cells per min for each chip with high delivery efficiency (>90%), cell viability (>80%), and uniform dosages (<60% coefficient of variation (CV)). In the current study, we modified the previous design into seven parallel microfluidic channels with the same number of of LCAT pairs (**Figure 2A-B**). We measured the cell traveling velocity within the acoustic microstreaming and derived the shear force that cells experienced (Supplementary Note). More specifically, cells exhibited a faster velocity near the air-liquid interface, compared to the velocity of the cells when they were away from the air-liquid interface. As a result, the acoustically activated microbubble generates microstreaming and exerts mechanical shear that deforms the cells orbiting in these microvortices. In Figure 2C, the cells experienced the highest amount of shear (~25 Pa) at location b near the air-liquid interface. When the cells moved away from the air-liquid interface, they experienced lower shear force. As the applied voltage increased from 2 to 6 V, the microbubble oscillation increased and led to an increased microstreaming velocity. The shear stress cells experienced at location B increased from 5 to 25 Pa. Here we further explored the relationship between the microfluidic channel height and throughput within the AESOP device; as the height increased from 40 to 60 μm, the number of processed cells also increased (Figure 2D). At the 6 Vpp condition, throughput increased from 1.8 M at a height of 40 μm to 3.2 M per min at a height of 60 μm.

**Figure 2.**
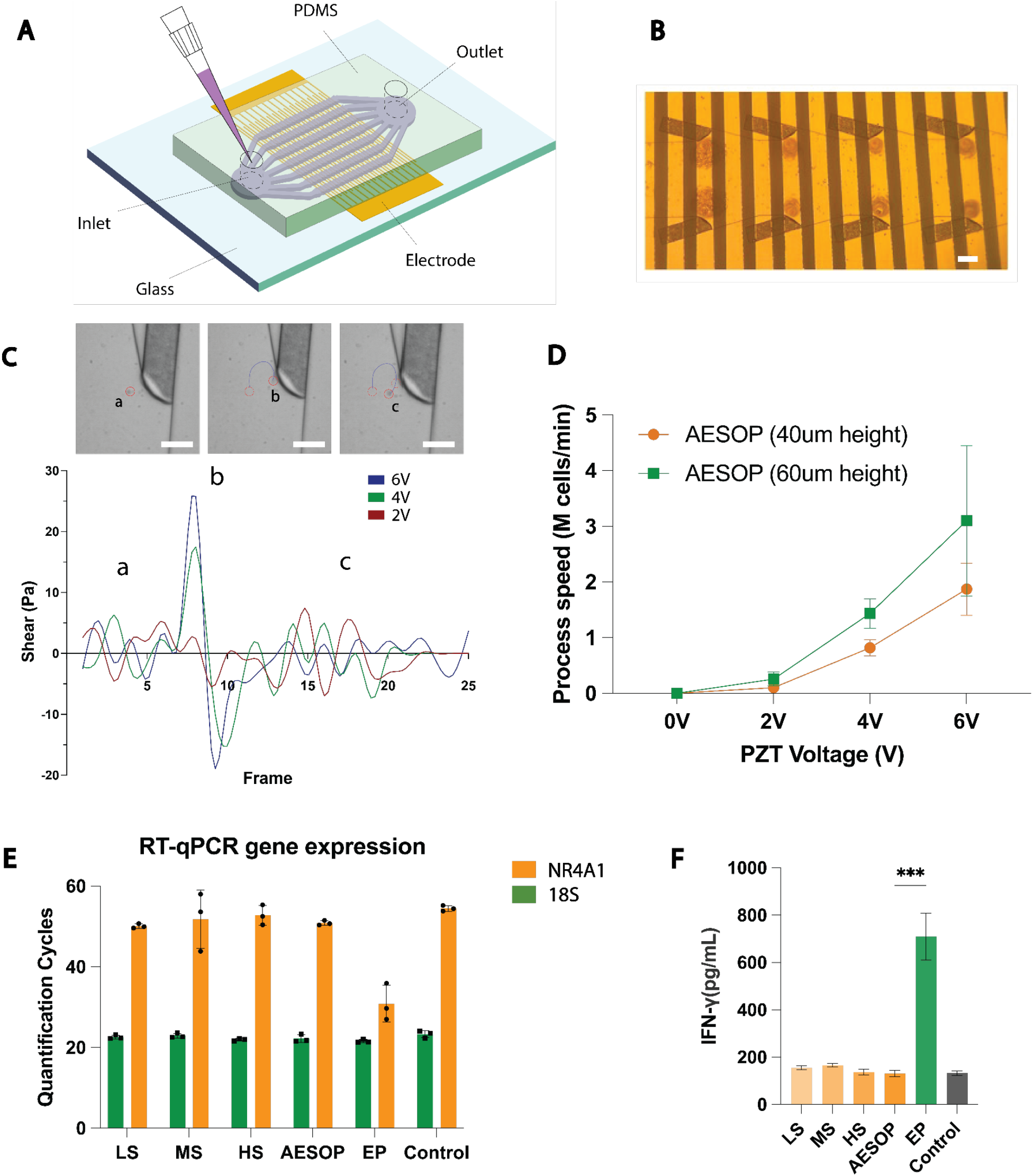
Characterization of AESOP. (A) Schematic diagram of the parallel AESOP platform. (B) Microscopic image of cells trapped in acoustic microstreaming vortices. The scale bar is 100 μm. (C) Cells experience shear measurements in vortices under PZT operation voltages of 2, 4, and 6 V over time. The scale bar is 100 μm. (D) The process speed and throughput (million cells/min) of the parallel AESOP platform under PZT operation voltages of 2, 4, and 6 V PZT with 40 μm and 60 μm channel heights. (E) Quantified human 18S and NR4A1 mRNA expression levels in T cells after electroporation and AESOP processing, which include low shear (2 V PZT voltage), moderate shear (4 V PZT voltage), and high shear (6 V PZT voltage) and AESOP (moderate shear and electric-pulse triggered). (F) T cell INF-γ measurement by ELISA after electroporation and AESOP processing.

### 2.2. Uniform electric field enlargement of shear-induced pores for cargo delivery

We investigated the performance of AESOP in intracellular delivery and transfection of mRNA. First, 500 kDa dextran, which the same size as mRNA, was mixed with T-cell pellets. Then, the cell solution was introduced into the microchannels to prime the device and form an air-liquid interface. The AESOP operation was then triggered by applying a PZT voltage and electric field. After 1–2 mins of cell mixing through acoustic microstreaming, T cells were collected and seeded into a 48-well culture plate. Delivery efficiency was evaluated 2 h after delivery using flow cytometry. The delivery efficiency in the AESOP group was 76% ± 9.76%, while the delivery efficiency in the electroporation group was 47.5% ± 12.74%, as shown in Figure 3A. In addition, the cell viability in the AESOP group was 80.4% ± 5.48%, compared to the 44.93% ± 8.29% cell viability in the electroporation group. We further delivered eGFP mRNA into primary T cells and measured the transfection efficiency for the GFP protein expression 24 h after the mRNA was delivered into the nucleus. The results showed that AESOP achieved > 70% transfection efficiency, with > 70% of the cells viable. In contrast, electroporation groups showed a transfection efficiency and cell viability of both approximately 50%.

**Figure 3.**
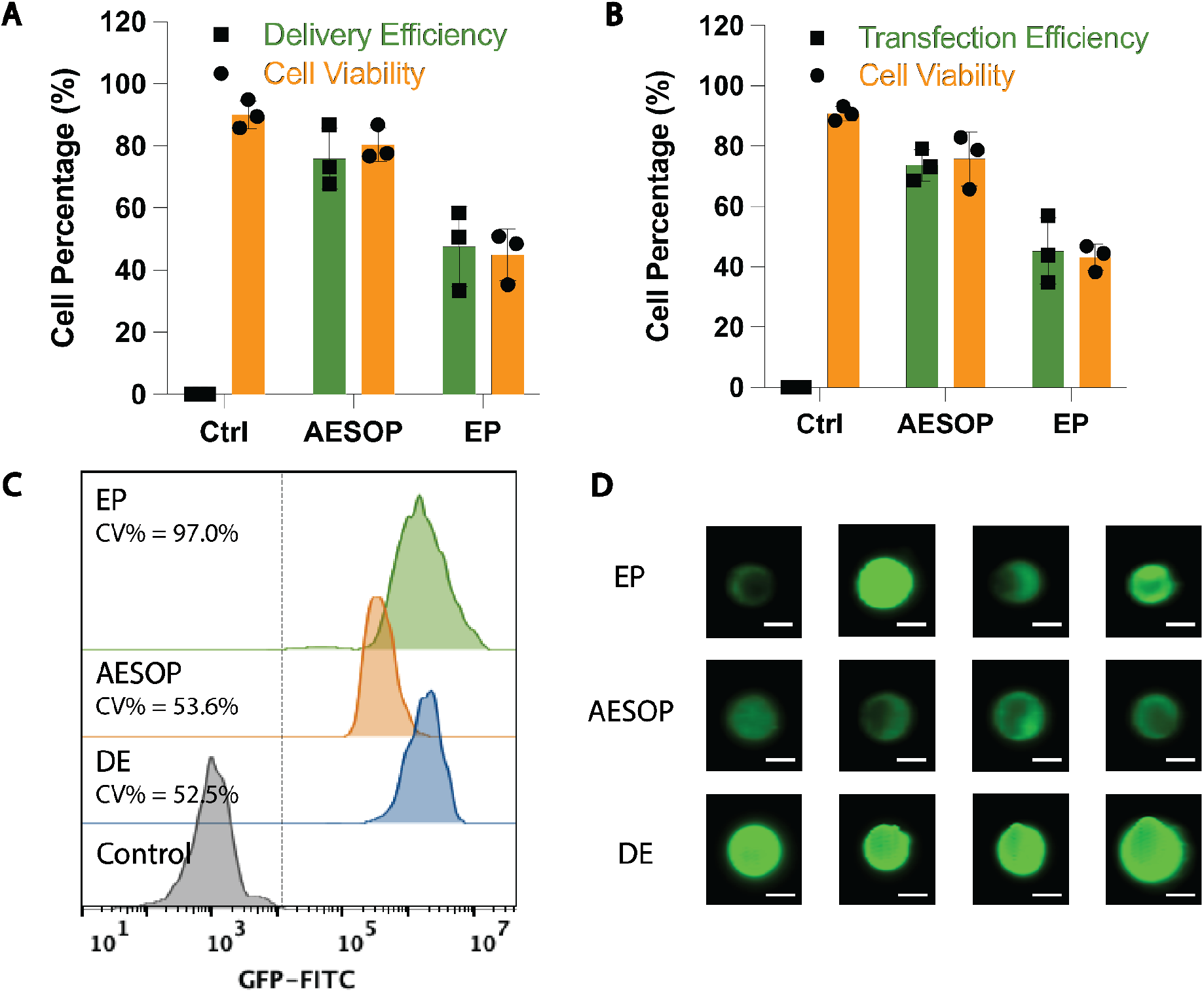
Efficiency and uniformity of AESOP for delivering 500 kDa dextran and eGFP mRNA. (A) Delivery efficiency and cell viability results measured 2 h after the delivery of 500 kDa dextran. (B) Transfection efficiency and cell viability results measured 24 h after the delivery of eGFP mRNA. (C) %CV to quantify the uniformity of delivery between AESOP, electroporation, and double emulsion (DE) groups. All the histograms were normalized for better comparison between each group. (D) Single-cell image after the delivery of 500 kDa dextran among electroporation, AESOP, double emulsion (DE) groups. The scale bar in all images is 7 μm.

In the AESOP platform, acoustic microstreaming vortices played a crucial role in uniformly mixing cells and cargoes and delivering a similar amount of cargo into each cell. 500 kDa dextran was used to quantify the uniformity of cellular uptake across the cell population. The percentage coefficient of variation (%CV, which is calculated as the percentage of the standard deviation to the mean) of the fluorescence intensity across the cell population was calculated to quantify uniformity. The lower the value of %CV, the more monodisperse the fluorescence intensity across the cell population, which indicates a more uniform dextran dosage taken by the cells. Conversely, the higher the value of %CV, the more varied the dextran dosage taken up by the cells. A cell-sized double emulsion (DE) encapsulating dextran solution as the inner phase, was used as a positive control group, which mimicked the situation where the cell uptake was completely uniform. Compared to the electroporation (EP) group, in which %CV was 97%, %CV of AESOP group was approximately 53.6% and was closer to that of DE group, with %CV of 52.5% (**Figure 3C**). Single-cell and DE images were acquired using a flow cytometer (**Figure 3D** and **S1**): DE and AESOP cell images showed the uniform fluorescence intensities within each group, whereas cell images from the electroporation group appeared to be non-uniform, with visible differences in terms of the fluorescence intensity. According to the results, AESOP performed intracellular delivery more uniformly than electroporation by reducing %CV from 97% to 53.6%.

### 2.3. Dosage-controlled capability and mechanism of intracellular delivery

In the previous section, we proved that the AESOP platform was able to deliver dextran into the cell population uniformly with the assistance of fluid mixing induced by acoustic microvortices, and the concentration of the delivered cargo could be further tuned. Therefore, we hypothesized that by adjusting the input concentration of reagents, the dosage required by each cell can be controlled because each cell will uptake the average amount of cargo from the input reagents. To test this hypothesis, we delivered various dextran concentrations (from 0.5 mg/mL, 1, 2, 4, and 8 mg/mL) into primary T cells using AESOP with 2 V PZT voltage, and 100 Vpp electric voltage, 30 kHz, and 200 cycles for electric pulses. Fluorescence intensity across the cell population was evaluated using flow cytometry. According to the histogram of fluorescence intensity, the fluorescence intensity peaks in the AESOP groups increased as the dextran concentration increased, whereas the fluorescence peaks in the electroporation groups did not increase once the dextran concentration was higher than 1 mg/mL (**Figure 4A**). As illustrated in **Fig. 4B**, the mean fluorescence intensity (MFI) was calculated from the flow cytometry histogram results for different dextran concentrations. Based on these results, the MFI, which was quantifies the amount of dextran delivered into cells, was more linear to the dextran concentration in the AESOP groups than in the electroporation groups. We then applied linear fitting to the MFI at various concentrations and the R-squared values for the AESOP and electroporation groups were 0.95 and 0.49, respectively, with the former indicating a higher fitness and suggesting that AESOP enables dosage-controlled delivery capabilities.

**Figure 4.**
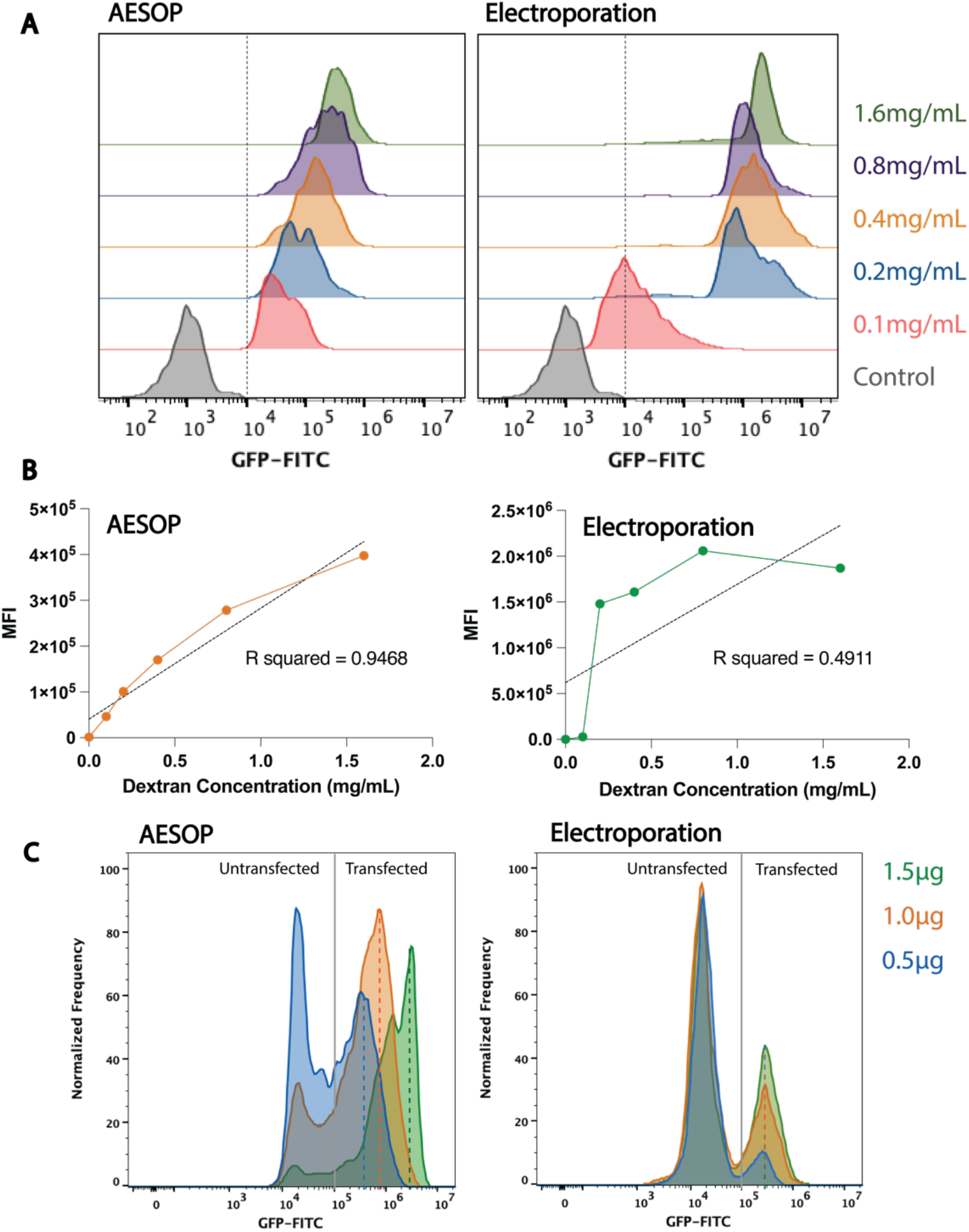
Dosage-controlled delivery and protein expression-level manipulation by AESOP. (A) Flow cytometry histogram of GFP fluorescence intensity of various input dextran concentrations. (B) Line graph showing the linear correlation between the mean fluorescence intensity (MFI) and dextran concentration. The R-squared values in AESOP and electroporation groups are 0.95 and 0.49, respectively. (C) Flow cytometry histogram of the GFP protein expression under different dosages of eGFP mRNA. The dash lines indicate the mean GFP protein expression level correspond with various dosages of mRNA.

In the next step, to further investigate the performance of controlled protein expression by AESOP which is positive correlated with mRNA abundance uptake by the cells,^[23]^ we introduced eGFP mRNA into primary T cells and measured the GFP expression level in the cell population using a flow cytometer. The dosage of eGFP mRNA (0.5 μg, 1.0, and 1.5 μg) was used for cell transfection. The results demonstrated that AESOP was able to control the expression levels of GFP protein by observing an increasing trend of fluorescence intensity as the mRNA dosage increased. The increasing trend was evidenced by the peaks of the fluorescence histogram shifting to the right. In contrast, for the electroporation groups although the transfection efficiency increased, the fluorescence histogram peaks remained at approximately the same value without a dynamic shift when the mRNA dosage increased (**Figure 4C**).

### 2.4. CAR mRNA transfection and CAR T-cell generation with dosage control

In the previous section, we proved that AESOP can deliver and control the dosage of cargoes, thus titering the protein expression level by eGFP mRNA as a proof of concept. In this section, CAR mRNA encoding anti-CD19 and GFP-reporter (**Figure S2**), which was manufactured from the pSLCAR-CD19-CD3z plasmid^[24]^ by in vitro transcription (IVT), is used to test CAR mDNA transfection. To verify whether the GFP reporter is a proper indicator of the CAR gene (anti-CD19), we measured the relationship between the GFP reporter and CAR expression. Transfected cells were labeled with anti-CD19 APC (**Figure 5C**). Linear correlation was observed between APC fluorescence intensity and GFP fluorescence intensity, indicating that the GFP reporter was a proper indicator of CAR expression (**Figure S3**). First, we explored the performance of AESOP for the intracellular delivery of CAR mRNA and obtained the transfection efficiency according to the GFP reporter 24 h after the experiment. According to the results, the AESOP group showed 64% transfection efficiency and 80% cell viability compared to the electroporation groups, which had transfection efficiency and cell viability both at approximately 35% (**Figure 5A-B**). Finally, the three different dosages of CAR-mRNA (2.5μg, 5.0μg, and 7.5μg) were delivered into primary T cells to investigate the corresponding CAR expression level. Based on these results, the APC fluorescence intensity, which corresponded to the CAR expression level on the cell membrane, increased when the dosage of CAR mRNA increased in AESOP. However, in the electroporation group, the CAR expression levels were widespread, and no fluorescence intensity shifts were observed as the CAR mRNA dosage increased.

**Figure 5.**
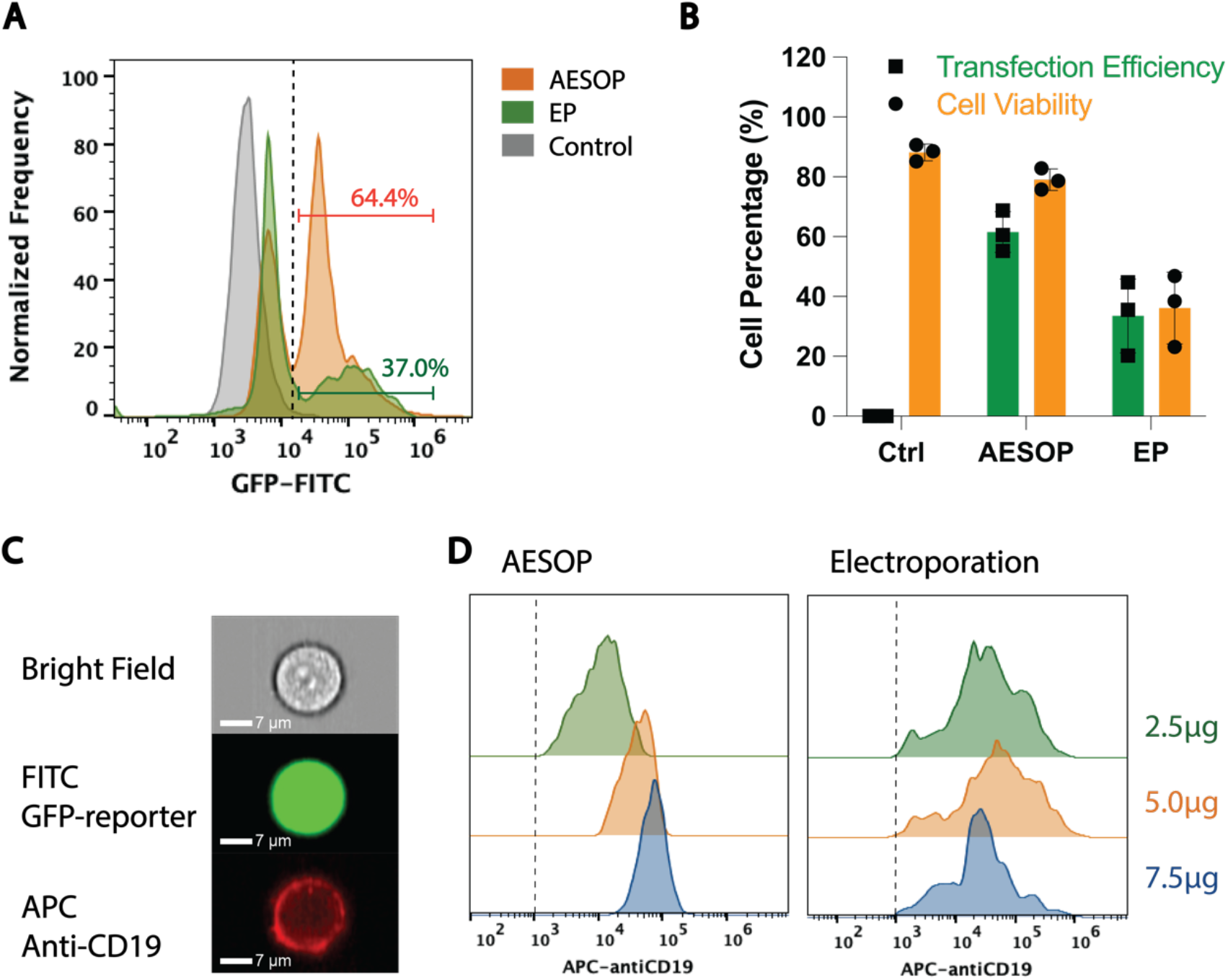
AESOP performance of CAR mRNA transfection and control of the CAR expression into the T cell population. (A) Flow cytometry histogram for quantifying CAR mRNA transfection efficiency according to GFP reporter. (B) Transfection efficiency of CAR mNRA and cell viability of T cells measured 24 h after intracellular delivery. (C) Magnified image of single CAR T cells with GFP reporter and label of anti-CD19 stained with an APC stain. (D) The histogram of fluorescence intensity used to measure anti-CD19 expression levels under various CAR mRNA dosages.

## 3. MATERIALS AND METHODS

### 3.1. Materials and Reagents

Fetal bovine serum (FBS), Iscove’s modified Dulbecco’s medium (IMDM), Dynabeads CD3/CD28, and recombinant human interleukin-2 (IL-2) protein (Invitrogen) were purchased from Thermo Fisher Scientific. K562 cells were purchased from the American Tissue Culture Collection (ATCC; Manassas, VA). ImmunoCult-XF T Cell Expansion Medium and immunomagnetic negative selection kits were purchased from STEMCELL Technologies. FITC-dextran molecules (500 kDa) was purchased from Millipore Sigma. CleanCap-Enhanced Green Fluorescent Protein mRNA was purchased from TriLink Biotechnologies. APC-Labeled Human CD19 Protein was purchased from Acro Biosystems.

### 3.2. AESOP Device Fabrication

AESOP integrates Interdigitated Array (IDA) electrodes with an LCAT microfluidic chip. For the IDA electrodes, the lift-off technique was used for fabrication. At the beginning of the liftoff process, the positive photoresist MICROPSIT S1813 was patterned with the outline of the electrode as a standard photolithography protocol. E-beam evaporation was performed to deposit 300 Å of chromium (Cr) followed by 1000 Å of gold (Au) on glass slides. Finally, the glass slides were sonicated in an acetone bath to remove the unwanted metal photoresist and the metal layer. For the microfluidic part, standard photolithography was used to fabricate the negative mold, in which a negative photoresist SU-8 2075 (Kayaku Advanced Materials, Inc.) was used for pattern fabrication on a silicon wafer. Finally, a polydimethylsiloxane (PDMS) (Sylgard 184, Dow Corning) base was mixed with a curing agent at a 10:1 ratio and poured onto the silicon wafer mold, degassed for 1h in a vacuum chamber, and cured at 65 °C overnight. Finally, the cured PDMS was bonded onto the glass slide with the electrode after the oxygen plasma treatment and baked overnight at 65 °C to allow the PDMS device to recover its hydrophobicity.

### 3.3. Primary T-cell isolation and culture protocols

Blood samples from healthy donors were obtained from the Institute for Clinical and Translational Science (ICTS), at the University of California, Irvine. Within 12 h of blood collection, primary T cells were isolated using immunomagnetic negative selection kits (STEMCELL Technologies) according to the suggested protocols. After isolation, the T cells were mixed with PBS-washed CD3/CD28 Dynabeads at a cell-to-bead ratio of 1:1. Isolated T cells and Dynabeads were cultured in the ImmunoCult-XF T Cell Expansion Medium with 30 UmL^-1^ human IL-2 recombinant protein at 37 °C in a humidified 5% CO_2_ incubator for 3 days at a seeding density of 1 × 10^6^ cells mL^-1^.

### 3.4. CAR mRNA Synthesis

pSLCAR-CD19-CD3z was a gift from Scott McComb (Addgene plasmid # 135993; http://n2t.net/addgene:135993; RRID:Addgene_135993)^[24]^. In vitro transcription (IVT) was performed by TriLink Biotechnologies to synthesize mRNA encoding CAR-CD19.

### 3.5. Setup of the RT-qPCR

The genotype of human primary T cells was characterized after processing with AESOP and electroporation. RT-qPCR was performed and measured in human 18S and NR4A1 mRNA expression quantitatively 3 days after the process. The Quick-RNA Microprep Kit purchased from Zymo Research was used to lyse cells, extract RNA, and purify RNA according to the manufacturer’s protocol. NanoDrop 2000 Spectrometers were used to measure the concentration and purity of the RNA samples (260/280 > 2.0). Next, the iScript™ cDNA Synthesis Kit (Bio-Rad) was used to synthesize 500 ng RNA from each sample into cDNA products, and the thermocycler protocol was 25 °C for 5 min, 42 °C for 30 min, 85 °C for 5 min, and then 4 °C for 10 min. Finally, 20μL of RT-qPCR mix was prepared with 10μL SYBR™ Green PCR Master Mix (Thermo Fisher Scientific), 8μL cDNA products, and 1 μL each forward and reverse primer. A S1000 Thermal Cycler (Bio-Rad) was used with the following thermal cycling setup: 95 °C for 3 min, 95 °C for 10 s, then 55 °C for 30 s, which repeated for 40 cycles. The sequences of the forward and reverse primers for human 18S rRNA were 5’-ATTCGAACGTCTGCCCTATCAA-3’, and 5’-CGGGAGTGGGTAATTTGCG-3’, respectively. The sequences of the forward and reverse primers for NR4A1 were 5’-ATGCCTCCCCTACCAATCTTC-3’, and 5’-CACCAGTTCCTGGAACTTGGA-3’, respectively. All the primers were purchased from Integrated DNA Technologies.

### 3.6. Flow cytometry analysis

After transfection with AESOP and eGFP or CAR mRNA, the cells were incubated for 24 h. The cells were washed and resuspended in a flow cytometry buffer (2% FBS in PBS). For the cell viability assay, Calcein Red AM (AAT Bioquest) was added to the cell solution at a 1:100 volume ratio to stain live cells. For CAR T-cell transfection, T cells were stained with APC-Labeled Human CD19 protein (Acro Biosystems), according to the manufacturer’s protocol.

The cell samples were scanned using an ImageStream Mark II Imaging Flow Cytometer (Amnis Corporation) at 60× magnification. Cell viability, transfection efficiency, and fluorescence intensity (%CV) were analyzed using IDEAS, a simulation software, (Amnis Corporation) and FlowJo.

### 3.7. T cells IFN-γ release measured by ELISA kit

IFN-γ release by T cells was quantified using ELISA MAX™ Standard Set Human IFN-γ, which was purchased from BioLegend. All reagents that were not included in the kit were purchased from BioLegend, according to the manufacturer’s instructions. Standard curves and samples were prepared according to the manufacturer’s instructions. A BioTek Cytation5 plate reader was used to acquire results using a colorimetric mold.

### 3.8. Electroporation transfection experiment

All electroporation experiments were performed using Lonza Nucleofector Transfection 2b Device. The electroporation buffer was obtained from the Human T Cell Nucleofector™ Kit and prepared according to the manufacturer’s protocol. The electroporation program used was T-020, according to the manufacturer’s instructions.

### 3.9. Double emulsion encapsulates dextran preparation

The double emulsion droplets (DEDs) were generated using a flow-focusing microfluidic channel. The inner phase was composed of 250 mM sucrose with 2 mg/ml FITC (fluorescein isothiocyanate). The oil phase was made of 7.5 mg/ml DOPC, 2.5 mg/ml DPPC, and 5 mg/ml cholesterol dissolved in oleic acid. A detailed droplet generation method and channel geometry can be found in our former study.^[25]^ In brief, the inner phase and oil phase were sheared by an external phase (15% glycerol + 125mM NaCl + 6% Pluronic F68) and droplets were pinched off. The DEDs were diluted in 1XPBS before sending to the flow cytometry.

#### 3.9.1. Image data processing and analysis

High-speed images were recorded using a phantom high-speed camera (Vision Research, USA). The cell traveling distance was measured using ImageJ, an analysis software, (https://imagej.nih.gov/ij/).

## 4. DISCUSSION

The antitumor efficacy of CAR T cells relies on the interactions between the receptors on engineered T cells and the ligands on the tumor cells. CAR is usually expressed by introducing tumor-specific gene sequences against tumor antigens such as anti-CD19 into the nucleus of T cells. The density of CAR molecules on the membrane of CAR T cells influence their heterogeneity, thereby affecting their functionalities and antitumor efficacies.^[26],[27]^ More specifically, CAR^High^ T cells are associated with excessive cytokine release, whereas CAR^Low^ T cells are not equipped with effective ligands that interact with tumor cells. The traditional non-viral approach used to introduce CAR genes is bulk electroporation, which operates on the principle of an electrical field opening and expanding nanopores on the cell membrane^[19]^. However, bulk electroporation applies an undesired strong electrical field to induce the initiation of membrane pores, which damages cell viabilities.^[28]^ Moreover, to date there are no reported platforms capable of controlling CAR expressions on the surface of T-cell membranes to produce homogeneous CAR T cells.

In previous studies, we have shown that AESOP permeabilizes the cell membrane and precisely controls cellular dosage uptake in an efficient, precise, and high-throughput manner. AESOP consists of a two-step membrane disruption strategy: mechanical shear pore initialization and electrical field pore modulation. First, AESOP employs LCAT’s bubble oscillation and microstreaming vortices to apply gentle, tunable, and uniform mechanical shear on cells near oscillating air-liquid interfaces to uniformly create nanopores on their membranes. Second, these nanopores are enlarged upon uniform exposure of the cells to gentle low-strength electric fields. After nanopore creation and expansion on cell membranes, acoustic microstreaming generates chaotic mixing to ensure that the cargo can be uniformly and efficiently delivered into the cells. Cells continuously mix with cargoes in acoustic microstreaming vortices, resulting in a uniform cellular uptake across the cell population.

To adopt CAR T-cell therapy in clinical settings, it is critical to satisfy the requirements for high-throughput cell processing.^[29]^ On average, many CAR T cells, ranging from millions to billions, are sufficient to eradicate tumors for successful treatment outcomes and patient survival. In this study, we modified our AESOP design to further increase the cell-processing density and speed from the previously reported 1 million cells per minute per chip to 3 million cells per minute per chip. Specifically, seven parallel microchannels with one shared inlet and one outlet intensified the bulk fluid containing T cells and mRNA cargoes to facilitate sample pumping/processing. Increasing the channel height from 40 to 60 μm further enhanced throughput from 1.5 million cells per minute for each chip to 3 million cells per minute for each chip.

Our group previously demonstrated that AESOP enabled the delivery of 6.1 kbp eGFP and 9.3 kbp CRISPR-Cas9 plasmids with high efficiency and viability. While the study provided a proof of concept that plasmid encoding CAR genes, with a size of approximately 9 kbp^[24]^ delivered into T cells using AESOP is possible, delivery using plasmid for CAR T-cell generation has several limitations: Plasmids need to enter the cell nucleus, and then become transcribed into mRNA. Once mRNA is formed, the mRNA molecule is released into the cytoplasm and translated into a protein. Consequently, plasmid delivery is slower than direct mRNA delivery, and poses delayed manufacturing challenges. More importantly, CD19-directed CAR T cells target both cancerous and normal B cells, and the elimination of all CD19 positive cells potentially results in B-cell aplasia^[30]^. Since plasmid-delivered CAR T cells exhibits permanent CAR expression. the integration of the new genome into the host nucleus presents several safety concerns owing to the insertion of foreign genetic materials and immunogenicity. Furthermore, the size of the plasmid (5–9 kbp)^[24]^ is much larger than that of mRNA (300–500 base pairs)^[31]^ and is more challenging to deliver. In contrast to plasmid, mRNA allows for the transient expression of CAR as it is translated without genomic integration and has the potential to cause fewer on-target, off-tumor effects. Most synthetic mRNA molecules can be designed quickly and mass-produced in a cost-effective manner. Additionally, the amount of mRNA delivered to T cells affects the level of CAR expression in T cells, indicating that mRNA-based CAR expressions may offer a means to modulate the side effects associated with CAR T-cell therapy, such as cytokine release syndrome. In this study we have shown that AESOP precisely controls the amount of intracellular dextran expression under different input dextran doses from 0.5 μg to 1.5 μg while bulk electroporation consistently generates the same intracellular dextran expression under different input dextran doses from 0.5 μg to 1.5 μg. This highlights the ability of AESOP to control the amount of protein expression and generate homogeneous cells. We then further expanded the study to generate CAR T cells that expressed anti-CD19 using AESOP, and our results demonstrate a dosage controlled anti-CD19 expression (greater than 65%) from CAR^High^, CAR^Medium^ to CAR^Low^. This lends itself to more precise cell engineering applications, and, to the best of our knowledge, no other studies have demonstrated the ability to titer the CAR expression.

Furthermore, %CV was used as an indicator of the relative dispersion of the amount of mRNA delivered to the cell population and our study showed that %CV <60 can be achieved by AESOP. Compared to the commercial Lonza electroporation platform, AESOP also demonstrated an order of magnitude higher cell viability and transfection efficiency. These are particularly important for controlling uniform CAR expressions while reducing the cost associated with CAR T-cell generation.

In summary, AESOP provides a well-controlled process with a minimal benchtop space requirement that is capable of generating uniform CAR T cells for clinical applications. Moreover, our platform can be used to study the effect of CAR density on T-cell functionality and clinical responses in cancer treatment. Future work will involve testing the co-culture of CAR T cells with cancer cells and examining cytotoxicity, cell exhaustion, and cytokine release in a more physiological anticancer environment. AESOP can also be used to more precisely engineer a variety of cells, such as stem cells, dendritic cells, and macrophages. Finally, the comprehension of fluid mechanics and electroporation offers an important tool for identifying new physical approaches for controlled uniform protein expressions that can be used to develop new cell therapies.

## Supporting Information

Supporting Information is available from the Wiley Online Library or from the author.

## Acknowledgements

This work was supported by the National Institute of Health (NIH) (R01GM145987). The authors thank the Institute for Clinical & Translational Science (ICTS) in University of California, Irvine (UCI) for providing blood from healthy donors. The authors would like to thank Dr. Jui-Yi Chen, who provided double emulsions for the experiments.

## Supplementary Note: Shear calculation from high-speed video by image processing

We derived shear based on F = m × a, where F denotes the shear force, m denotes the mass of the individual T cell, and a is the acceleration of the T cell.

In contrast, F is equal to τ*A, where τ is the shear pressure, and A is the cross-sectional area of the individual T cell.

Making these two equations equal gives us:

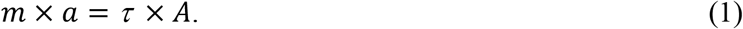

To rearrange the equation above:

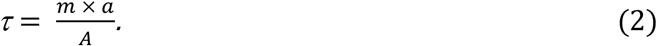

Acceleration (a) was calculated by dividing the velocity of cells’ movement over time. The mass of the T cells was taken from 3.5 ng.^[[32]]^

**Figure S1.**
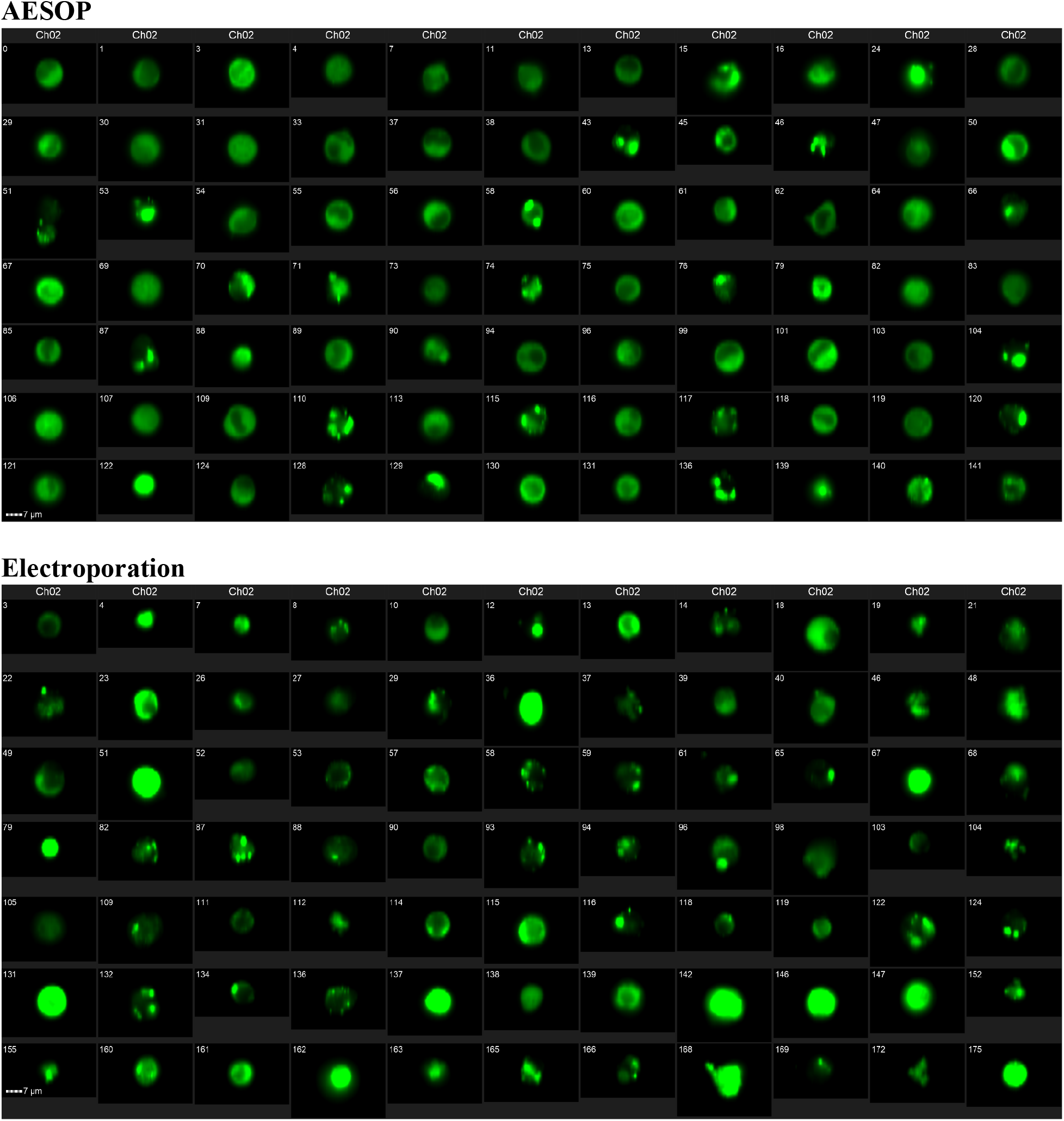
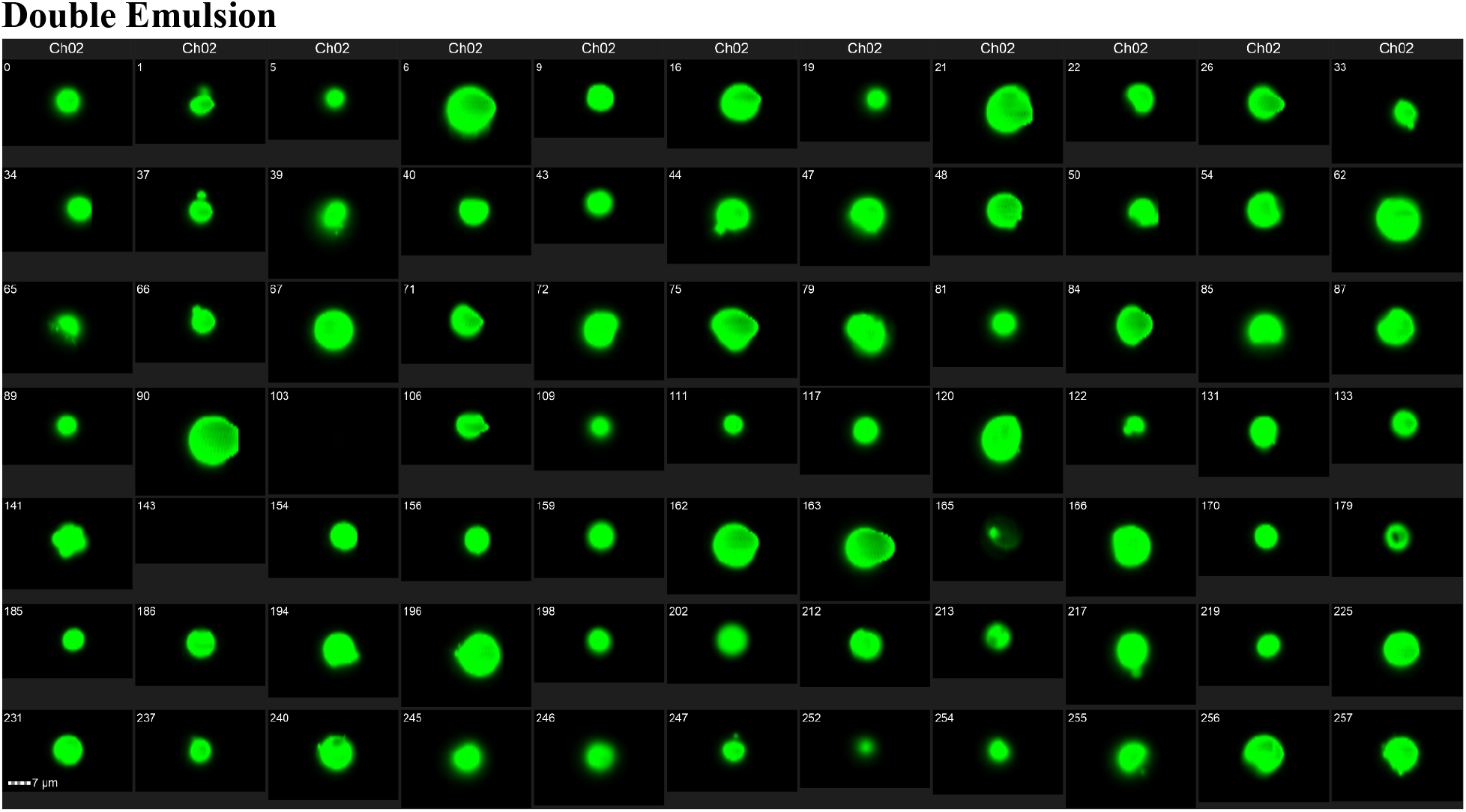
Single-cell image acquired from flow cytometers in AESOP, electroporation, and double emulsion groups to present the uniformity of fluorescence intensity in the cell/DE population.

**Figure S2.**
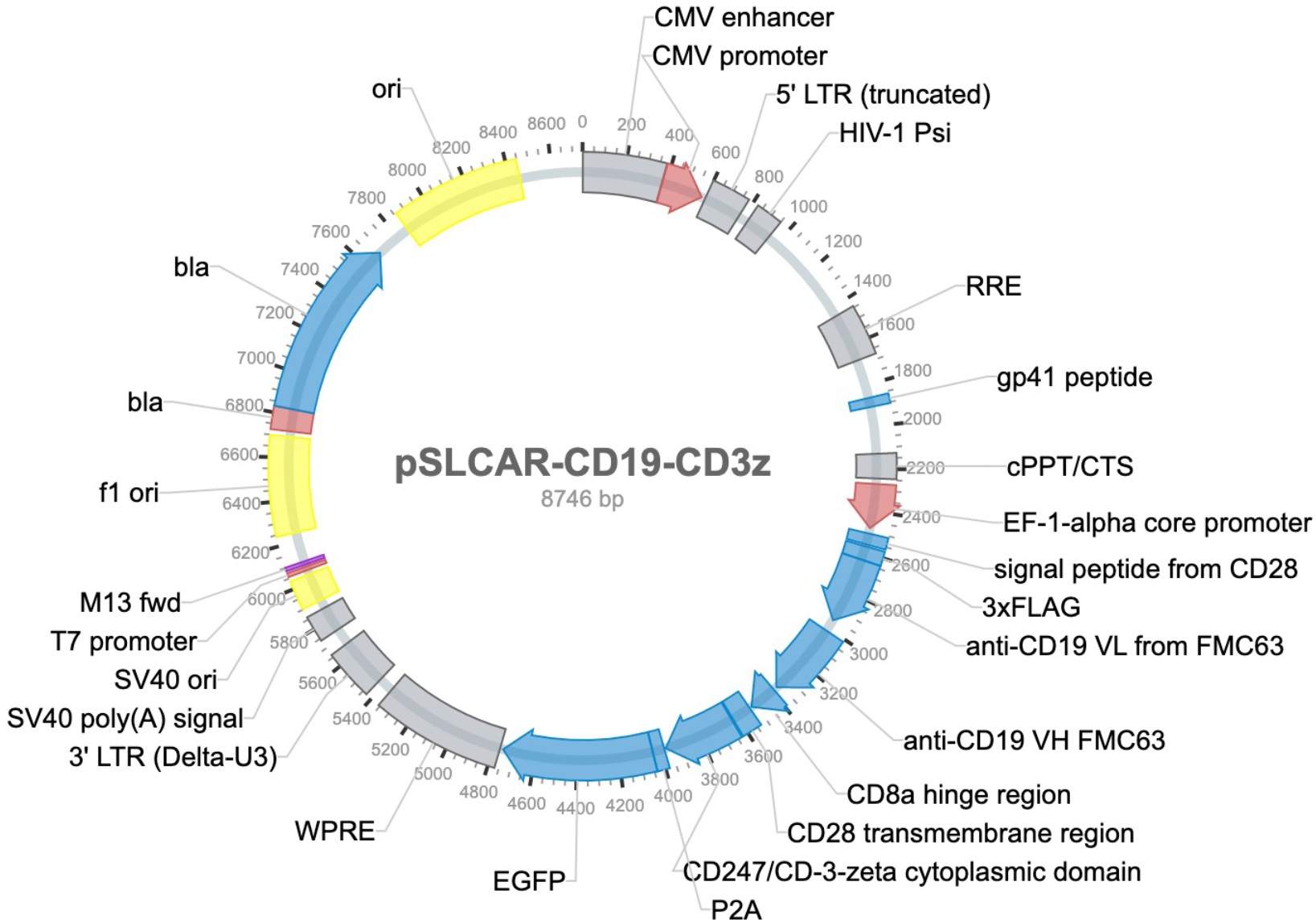
Plasmid map of pSLCAR-CD19-CD3z plasmid, which was a gift from Scott McComb (Addgene plasmid # 135993; http://n2t.net/addgene:135993; RRID:Addgene 135993).

**Figure S3.**
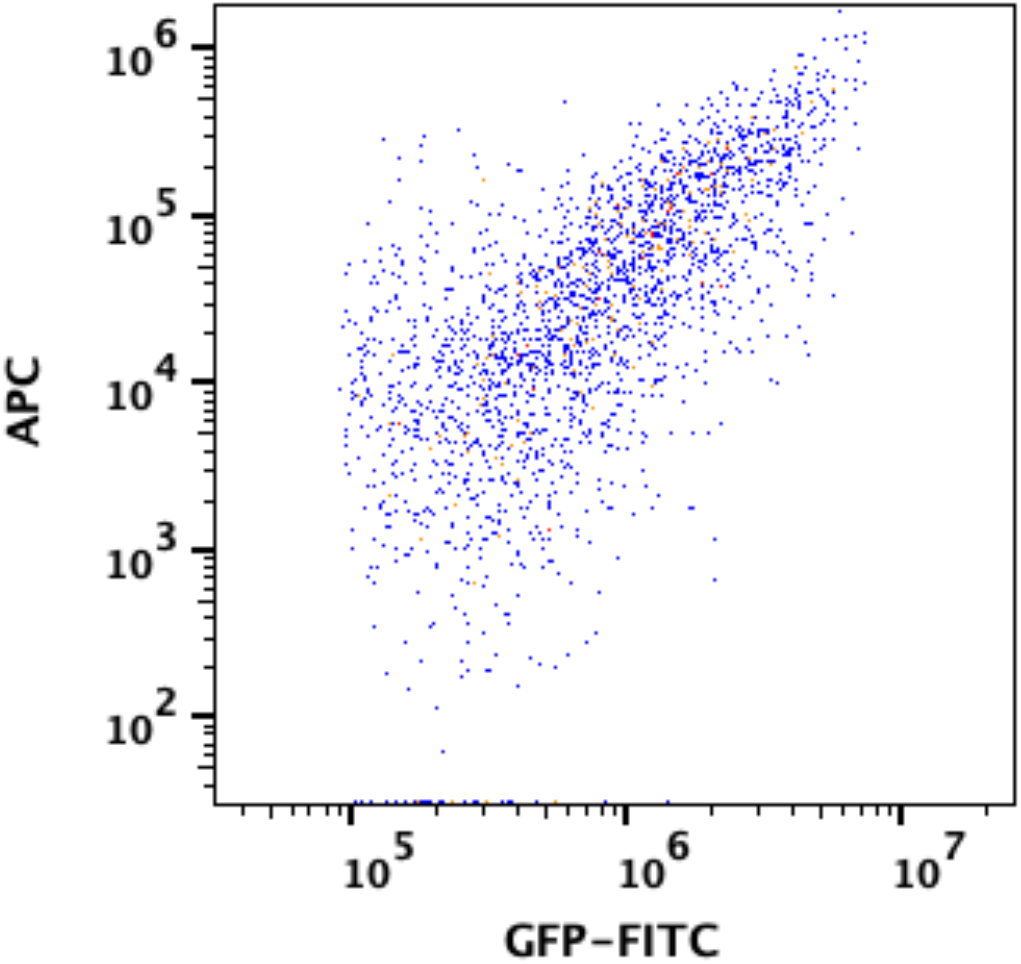
Correlation between the GFP reporter and APC-labeled anti-CD19 expressions in a CAR T-cell population.

## Notes

### Competing Interest Statement

The authors have declared no competing interest.

